# Repeated imitation makes human vocalizations more word-like

**DOI:** 10.1101/149708

**Authors:** Pierce Edmiston, Marcus Perlman, Gary Lupyan

## Abstract

People have long pondered the evolution of language and the origin of words. Here, we investigate how conventional spoken words might emerge from imitations of environmental sounds. Does the repeated imitation of an environmental sound gradually give rise to more word-like forms? In what ways do these forms resemble the original sounds that motivated them (i.e., exhibit iconicity)? Participants played a version of the children’s game “Telephone”. The first generation of participants imitated recognizable environmental sounds (e.g., glass breaking, water splashing). Subsequent generations imitated the previous generation of imitations for a maximum of 8 generations. The results showed that the imitations became more stable and word-like, and later imitations were easier to learn as category labels. At the same time, even after 8 generations, both spoken imitations and their written transcriptions could be matched above chance to the category of environmental sound that motivated them. These results show how repeated imitation can create progressively more word-like forms while continuing to retain a resemblance to the original sound that motivated them, and speak to the possible role of human vocal imitation in explaining the origins of at least some spoken words.

## Repeated imitation makes human vocalizations more word-like

Most vocal communication of non-human primates is based on species-typical calls that are highly similar across generations and between populations [1]. In contrast, human languages comprise a vast repertoire of learned meaningful elements (words and other morphemes) which can number in the tens of thousands or more [2]. Aside from their number, the words of different natural languages are characterized by their extreme diversity [3,4]. The words used within a speech community change relatively quickly over generations compared to the evolution of vocal signals [5]. At least in part as a consequence of this rapid change, most words appear to bear a largely arbitrary relationship between their form and their meaning — seemingly, a product of their idiosyncratic etymological histories [6,7]. The apparently arbitrary nature of spoken vocabularies presents a quandary for the study of language origins. If words of spoken languages are truly arbitrary, by what process were the first words ever coined?

While the origin of most spoken words remains opaque, the situation is somewhat different for signed languages for which much is known regarding the origins of many signs. Although signed languages rely on the same type of referential symbolism as spoken languages, many individual signs have clear iconic roots, formed from gestures that resemble their meaning [8–10]. For instance, [11] noted the iconic origins of the American Sign Language (ASL) sign for “bird”, which is formed with a beak-like handshape articulated in front of the nose. Another example is “steal”, derived from a grabbing motion to represent the act of stealing something. [12] identified about 25% of ASL signs to be iconic, and reviewing the remaining 75% of ASL signs, [13] determined that about two-thirds of these seemed plausibly derived from iconic origins. Further support for iconic origins of signed languages comes from observations of deaf children raised without exposure to a signed language, who develop homesign systems to use with their family. In these communication systems, children frequently use pantomimes and various iconic and indexical gestures some of which may become conventionalized [14]. Participants in laboratory experiments utilize a similar strategy when they cannot rely on existing words [15].

In contrast to the visual gestures of signed languages, many have argued that iconic vocalizations could not have played a significant role in the origin of spoken words because the vocal modality simply does not afford much form-meaning iconicity [16–21]. It has also been argued that the human capacity for vocal imitation is a domain-specific skill, geared towards learning to speak, rather than the representation of environmental sounds. For example, [22] suggested that, “most humans lack the ability. .. to convincingly reproduce environmental sounds. .. Thus ‘capacity for vocal imitation’ in humans might be better described as a capacity to learn to produce speech” (p. 209). Consequently, it is still widely assumed that vocal imitation — or more broadly, the use of any sort of resemblance between form and meaning — cannot be important to understanding the origin of spoken words.

Although most words of contemporary spoken languages are not clearly imitative in origin, there has been a growing recognition of the importance of iconicity in spoken languages [23,24] and the common use of vocal imitation and depiction in spoken discourse [25,26]. This has led some to argue for the importance of imitation for understanding the origin of spoken words [27–31]. In addition, counter to previous assumptions, people are highly effective at using vocal imitations to refer to events such as coins dropping in a jar or environmental sounds like scraping — even more effective in some cases than when using conventional words [32]. These imitations are effective not because people can mimic environmental sounds with high fidelity, but because people can capture with their “imitations” salient features of the referent in ways that are understandable to listeners [33]. Similarly, the features of onomatopoeic words might highlight distinctive aspects of the sounds they represent. For example, the initial voiced, plosive /b/ in “boom” represents an abrupt, loud onset, the back vowel /u/ a low pitch, and the nasalized /m/ a slow, muffled decay [34]. Such iconicity is not limited to imitations of sounds. People are able to create novel imitative vocalizations for more abstract meanings (e.g. “slow”, “rough”, “good”, “many”) such that the vocalizations are understandable to naïve listeners [31].

Thus, converging evidence suggests that people can use vocal imitation as an effective means of communication. At the same time, vocal imitations are not words. If vocal imitation played a role in the origin of some spoken words, then it is necessary to identify circumstances in which vocal imitation may begin can give rise to more word-like vocalizations that can eventually be integrated into a vocabulary of a language. In the present set of studies we ask whether vocal imitations can transition to more word-like forms through sheer repetition — without an explicit intent to communicate. To answer this question, we recruited participants to play an online version of the children’s game of “Telephone”. In our version of the game the original message (the “seed”) was a recording of an environmental sound. The initial group of participants imitated these seed sounds. The next generation imitated the previous imitators, and so on for up to 8 generations.

Our approach uses a transmission chain methodology similar to that frequently used in experimental studies of language evolution [35]. As with other transmission chain studies (and iterated learning studies more generally), we sought to discover how various biases and constraints of individuals changed the nature of a linguistic signal. While typical transmission chain studies focus on the impact of learning biases [36], here we use iterated reproduction which does not involve any learning. Participants simply attempt to imitate a sound as best as they can.

After collecting the imitations, we conducted a series of analyses and additional experiments to systematically answer the following questions: First, do imitations stabilize in form and become more word-like as they are repeated? Second, do the imitations retain a resemblance to the original environmental sound that inspired them? If so, it should be possible for naïve participants to match the emergent words back to the original seed sounds. Third, do the imitations become more suitable as categorical labels for the sounds that motivated them? For example, does the imitation of a particular water-splashing sound become, over generations of repeated imitation, a better label for the more general category of water-splashing sounds?

## Experiment 1: Stabilization of imitations through repetition

In the first experiment, we collected the vocal imitations, and assessed the extent to which repeating imitations of environmental sounds results in progressive stabilization toward more word-like forms in three ways. First, we measured changes in the perception of acoustic similarity between subsequent generations of imitations. Second, we used algorithmic measures of acoustic similarity to assess the similarity of imitations sampled within and between transmission chains. Third, we obtained transcriptions of imitations, and measured the extent to which later generation imitations were transcribed with greater consistency and agreement. The results show that repeated imitation results in vocalizations that are easier to repeat with high fidelity and more consistently transcribed into English letters.

## Methods

### Selecting seed sounds

To avoid sounds with lexicalized or conventionalized onomatopoeic forms in English, we used inanimate categories of environmental sounds. We ensured that the sounds within each category were approximately equally distinguishable by using an odd-one-out norming procedure (*N* = 105 participants; see Fig. S1), resulting in a final set of 16 sounds, 4 in each of 4 categories: glass (breaking), paper (tearing), water (splashing), zipper (moving).

### Collecting vocal imitations

We recruited 94 participants from Amazon Mechanical Turk. Participants were instructed that they would hear some sound and their task was to reproduce it as accurately as possible using their computer microphone. Full instructions are provided in the Supplemental Materials.

Each participant listened to and imitated four sounds: one from each of the four categories. Sounds were assigned at random such that participants were unlikely to imitate the same person more than once. Participants were allowed to listen to each target sound as many times as they wished, but were only allowed a single recording in response. Recordings that were too quiet (less than −30 dBFS) were not accepted.

A total of 115 (24%) imitations were removed for being poor quality (e.g., loud background sounds) or for violating the rules of the experiment (e.g., an utterance in English). The final sample contained 365 imitations along 105 contiguous transmission chains (Fig. 1).

**Figure 1.**
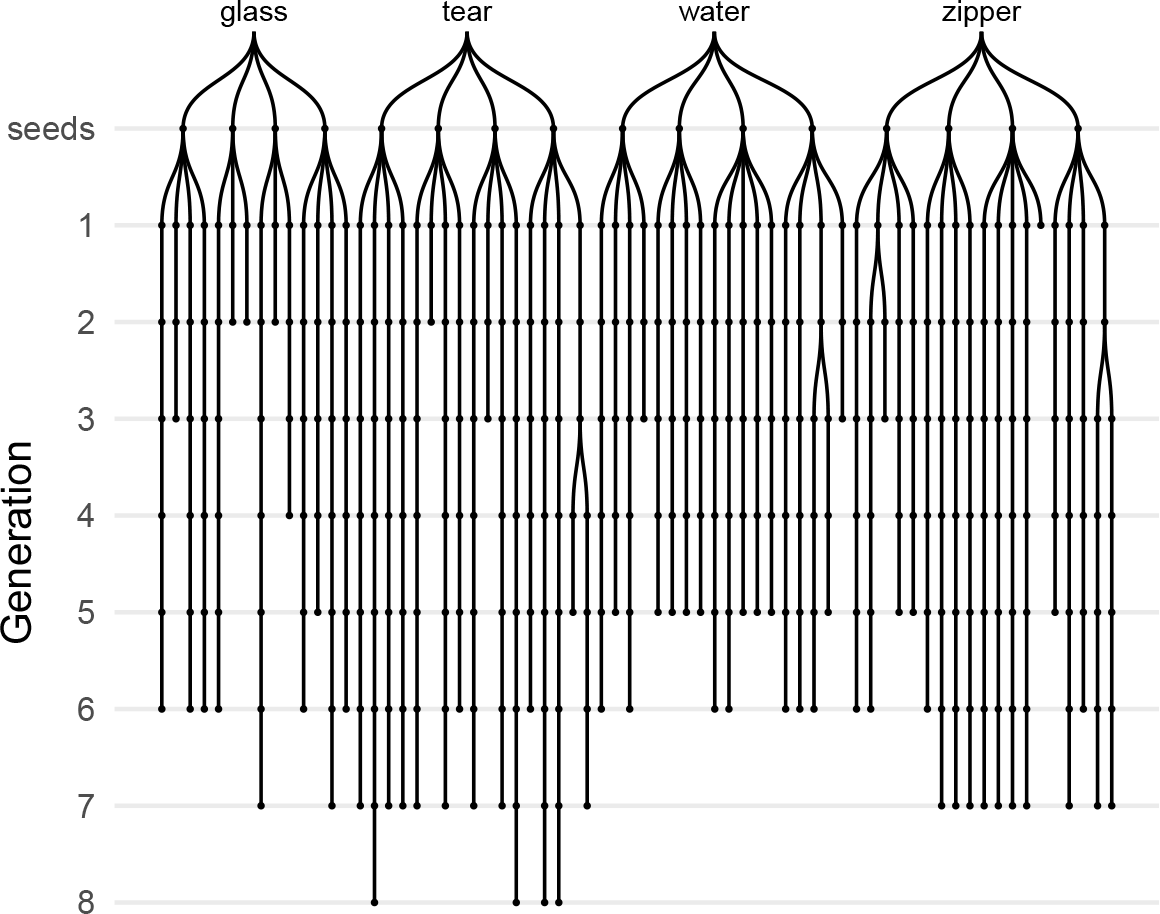
Vocal imitations collected in the transmission chain experiment. Seed sounds (16) were sampled from four categories of environmental sounds: glass, tear, water, zipper. Participants imitated each seed sound, and then the next generation of participants imitated the imitations, and so on, for up to 8 generations. Chains are unbalanced due to random assignment and the above-mentioned exclusion criteria.

### Measuring acoustic similarity

We obtained acoustic similarity judgments from five research assistants who listened to pairs of sounds (approx. 300 each) and rated their subjective similarity. On each trial, raters heard two sounds from subsequent generations played in random order, and indicated the similarity between the sounds on a 7-point Likert scale from *Entirely different and would never be confused* to *Nearly identical*. See Supplemental Materials for full instructions and inter-rater reliability measures.

We also obtained algorithmic measures of acoustic similarity using the acoustic distance functions from the Phonological Corpus Tools [37]. We computed Mel-frequency cepstral coefficients (MFCCs) between pairs of imitations using 12 coefficients in order to obtain speaker-independent estimates.

### Collecting transcriptions of imitations

Transcriptions were obtained for the first and last three generations of each transmission chain. We also transcribed the original seed sounds(see Supplementary Materials, Fig. S6).

We recruited 216 additional participants from Amazon Mechanical Turk to listen to the vocal imitations and write down what they heard as a single “word” so that the written word would sound as much like the sound as possible. Participants were instructed to avoid using English words in their transcriptions. Each participant completed 10 transcriptions.

## Results

Imitations of environmental sounds became more stable over the course of being repeated as revealed by increasing acoustic similarity judgments along individual transmission chains. Acoustic similarity ratings were fit with a linear mixed-effects model predicting perceived acoustic similarity from generation with random effects (intercepts and slopes) for raters. To test whether the hypothesized increase in acoustic similarity was true across all seed sounds and categories, we added random effects (intercepts and slopes) for seed sounds nested within categories. The results showed that, across raters and seeds, imitations from later generations were rated as sounding more similar to one another than imitations from earlier generations, *b* = 0.10 (SE = 0.03), *t*(11.9) = 3.03, *p* = 0.011 (Fig. 2). This result suggests that imitations became more stable (i.e., easier to imitate with high fidelity) with each generation of repetition.

**Figure 2.**
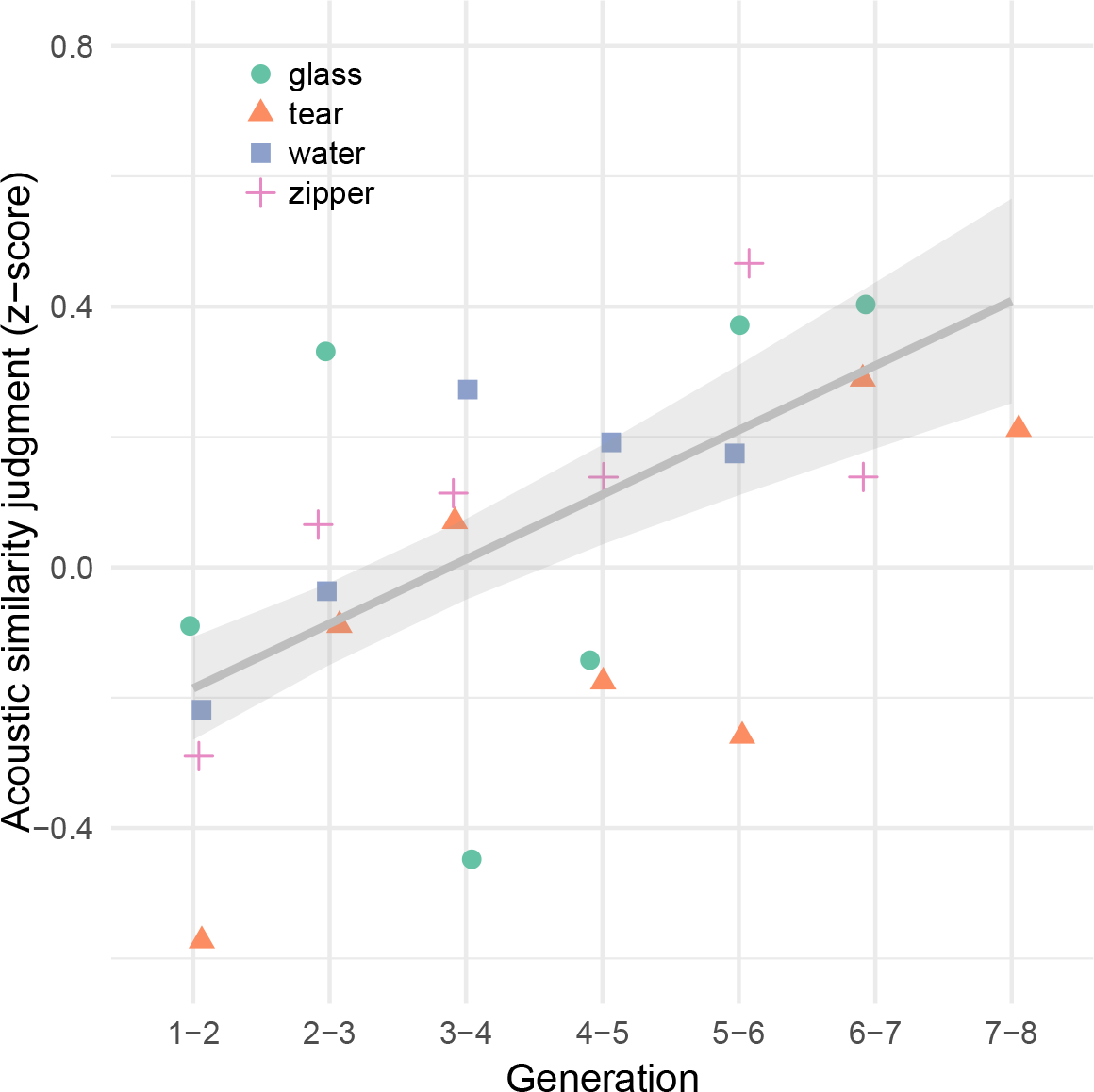
Change in perception of acoustic similarity over generations of iterated imitation. Points depict mean acoustic similarity ratings for pairs of imitations in each category. The predictions of the linear mixed-effects model are shown with ±1 SE.

Although in some chains, imitations were repeated up to 8 times, an increase in similarity between generations could be detected after about 5 generations. Imitations from chains that did not reach 5 generations due to experimental constraints (see Fig. 1) were included in all analyses, which included appropriate random effects to ensure that shorter chains were weighed appropriately in the analyses. However, chains with fewer than 5 generations were excluded from analyses involving transcriptions of the first and last imitation in each chain because these analyses collapse across generation.

Increasing similarity along transmission chains could also reflect the uniform degradation of the signal due to repeated imitation, in which case acoustic similarity would increase both within as well as between chains. To test this, we calculated MFCCs for pairs of sounds sampled from within and between transmission chains across categories, and fit a linear model predicting acoustic similarity from the generation of sounds. We found that acoustic similarity increased within chains more than it increased between chains, *b* = −0.07 (SE = 0.03), *t*(6674.0) = −2.13, *p* = 0.033 (Fig. S2), indicating that imitations were stabilizing on divergent acoustic forms as opposed to converging on similar forms through continuous degradation.

As an additional test of stabilization we measured whether later generation imitations were transcribed more consistently than first generation imitations. We collected a total of 2163 transcriptions — approximately 20 transcriptions per sound. Of these, 179 transcriptions (8%) were removed because they contained English words. Some examples of the final transcriptions are presented in Table 1.

**Table 1.** Examples of words transcribed from imitations.

| Category | First generation | Last generation |
| --- | --- | --- |
| glass | dirrng | wayew |
| tear | feeshefee | cheecheea |
| water | boococucuwich | galong |
| zipper | bzzzzup | izzip |

To measure the similarity among transcriptions for a given imitation, we calculated the average orthographic distance between the most frequent transcription and all other transcriptions of the same imitation. We then fit a hierarchical linear model predicting orthographic distance from the generation of the imitation (First generation, Last generation) with random effects (intercepts and slopes) for seed sound nested within category. The results showed that transcriptions of last generation imitations were more similar to one another than transcriptions of first generation imitations, *b* = −0.12 (SE = 0.03), *t*(3.0) = −3.62, *p* = 0.035 (Fig. S3). The same result is reached through alternative measures of orthographic distance (Fig. S4). Differences between transcriptions of human vocalizations and transcriptions directly of environmental sound cues are reported in the Supplementary Materials (Fig. S6).

## Discussion

Repeating imitations of environmental sounds over generations of imitators was sufficient to create more word-like forms (defined here in terms of acoustic stability and orthographic agreement), even without any explicit intent to communicate. With each repetition, the acoustic forms of the imitations became more similar to one another, indicating that it became easier to repeat them with greater consistency. The possibility that this similarity was due to uniform degradation across all transmission chains was ruled out by algorithmic analyses of acoustic similarity demonstrating that acoustic similarity increased within chains but not between them. Further support for our hypothesis that repeating imitations makes them more stable/word-like comes from the result showing that later generation imitations were transcribed more consistently into English letters.

The results of Experiment 1 demonstrate the ease with which iterated imitation gives rise to more stable forms. However, the results do not address how these emergent words relate to the original sounds that were being imitated. As the imitations became more stable, were they stabilizing on arbitrary acoustic and orthographic forms, or did they maintain some resemblance to the environmental sounds that motivated them? The purpose of Experiment 2 was to assess the extent to which repeated imitations and their transcriptions maintained a resemblance to the original set of seed sounds.

## Experiment 2: Resemblance of imitations to original seed sounds

To assess the resemblance of repeated imitations to the original seed sounds, we measured the ability of naïve participants to match imitations and their transcriptions back to their original sound source relative to other seed sounds from either the same category or from different categories (Fig. 3A). Using these match accuracies, we first asked whether and for how many generations the imitations and their transcriptions could be matched back to the original sounds and whether certain types of information were lost fater than other types. Specifically, we tested the hypothesis that if imitations were becoming more word-like, then they should also be interpreted more categorically, and thus we anticipated that imitations would lose information identifying the specific source of an imitation more rapidly than category information that identifies the category of environmental sound being imitated.

**Figure 3.**
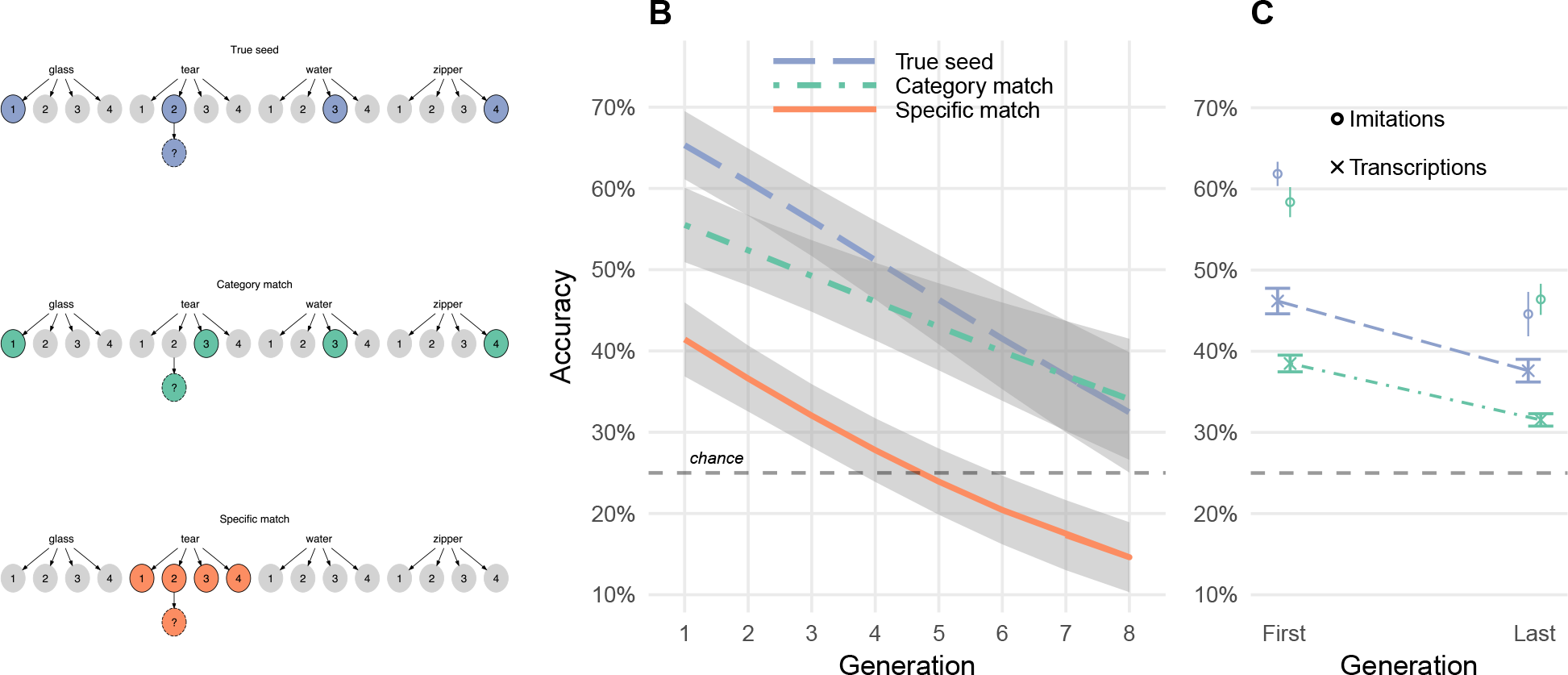
Repeated imitations retained category resemblance. A. Three types of matching questions. True seed and category match questions contained choices from different sound categories. Specific match questions pitted the actual seed against the other seeds within the same category. B. Accuracy in matching vocal imitations to original seed sounds. Curves show predictions of the generalized linear mixed effects models with ±1 SE of the model predictions. C. Accuracy in matching transcriptions of the imitations to original seed sounds (e.g., “boococucuwich” to a water splashing sound). Circles show mean matching accuracy for the vocal imitations that were transcribed for comparison.

### Methods

#### Matching imitations to seed sounds

Participants (*N*=751) recruited from Amazon Mechanical Turk were paid to listen to imitations, one at a time, and for each one, choose one of four possible sounds they thought the person was trying to imitate. The task was not speeded and no feedback was provided. Participants completed 10 questions at a time.

All imitations were tested in three question types (True seed, Category match, Specific match) which differed in the relationship between the imitation and the four seed sounds provided as the choices in the question (see Fig. 3A). The Question types were assigned between-subject.

#### Matching transcriptions to seed sounds

We recruited *N*=461 participants from Amazon Mechanical Turk to complete a modified version of the matching survey described above. Instead of listening to imitations, participants now saw a transcription of an imitation and were told that it was invented to describe one of the four presented sounds. Of the unique transcriptions that were generated for each sound (imitations and seed sounds), only the top four most frequent transcriptions were used in the matching experiment. The distractors for all questions were between-category, i.e. true seed and category match. Specific match questions were omitted.

### Results

Response accuracies in matching imitations to seed sounds were fit by a generalized linear mixed-effects model predicting match accuracy as different from chance (25%) based on the type of question being answered (True seed, Category match, Specific match) and the generation of the imitation. Question types were contrast coded using Category match questions as the baseline condition in comparison to the other two question types, each containing the actual seed that generated the imitation as one of the choices. The model included random intercepts for participant, and random slopes and intercepts for seed sounds nested within categories.

Accuracy in matching first generation imitations to seed sounds was above chance for all question types, *b* = 1.65 (SE = 0.14) log-odds, odds = 0.50, *z* = 11.58, *p* < 0.001, and decreased steadily over generations, *b* = −0.16 (SE = 0.04) log-odds, *z* = −3.72, *p* < 0.001. After 8 generations, imitations were still recognizable, *b* = 0.55 (SE = 0.30) log-odds, odds = −0.59, *z* = 1.87, *p* = 0.062. We then tested whether this increase in difficulty was constant across the three types of questions. The results are shown in Fig. 3B. Performance decreased over generations more rapidly for specific match questions that required a within-category distinction than for category match questions that required a between-category distinction, *b* = −0.08 (SE = 0.03) log-odds, *z* = −2.68, *p* = 0.007. This suggests that the iconicity in between-category information was more resistant to loss through repetition.

An alternative explanation of the relatively greater decrease in accuracy for specific match questions is that they are simply more difficult than the category-match questions because the sounds presented as choices are more acoustically similar to one another. However, performance also decreased relative to the category match questions for the easiest type of question where the correct answer was the actual seed generating the imitation (True seed questions; see Fig. 3A). That is, the advantage of having the true seed among between-category distractors decreased over generations, *b* = −0.07 (SE = 0.02) log-odds, *z* = −2.77, *p* = 0.006. Together, the observed decrease in the “true seed advantage” (the advantage of having the actual seed among the choices) and the increase in the “category advantage” (the advantage of having between-category distractors) shows that the changes induced by repeated imitation caused the imitations to lose some of properties that linked the earlier imitations to the specific sound that motivated them, while nevertheless preserving a more abstract category-based resemblance.

We next report the results of matching the written transcriptions of the auditory sounds back to the original environmental sounds. Remarkably, participants were able to guess the correct meaning of a word that was transcribed from an imitation that had been repeated up to 8 times, *b* = 0.83 (SE = 0.13) log-odds, odds = −0.18, *z* = 6.46, *p* < 0.001 (Fig. 3C) both for True seed questions containing the actual seed generating the transcribed imitation, *b* = 0.75 (SE = 0.15) log-odds, *z* = 4.87, *p* < 0.001, and for Category match questions where participants had to associate transcriptions with a particular category of environmental sounds, *b* = 1.02 (SE = 0.16) log-odds, *z* = 6.39, *p* < 0.001. The effect of generation did not vary across these question types, *b* = 0.05 (SE = 0.10) log-odds, *z* = 0.47, *p* = 0.638. The results of matching “transcriptions” directly of the environmental sounds are shown in Fig. S6.

### Discussion

Even after being repeated up to 8 times across 8 different individuals, vocalizations retained a resemblance to the environmental sound that motivated them. This resemblance remained even after the vocalizations were transcribed into orthographic forms. For vocal imitations, but not for transcriptions, this resemblance was stronger for the category of environmental sound than the specific seed sound, suggesting that iterated imitation produces vocalizations that are interpreted by naïve listeners in a more categorical way. Iterated imitation appears to strip the vocalizations of some of the characteristics that individuate each particular sound while maintaining some category-based resemblance. This happenned even though participants were never informed about the meaning of the vocalizations and were not trying to communicate.

Transcriptions of the vocalizations, like the vocalizations themselves, were able to be matched to the original environmental sounds at levels above chance. Unlike vocalizations, the transcriptions continued to be matched more accurately to the true seed compared to the general category; transcription appearred to impact specific and category-level information equally. One possible explanation of the difference between the acoustic and orthographic forms of this task is that the process of transcribing a non-linguistic vocalization into a written word encourages transcribers to emphasize individuating information about the vocalization. However, this does not provide a complete explanation of our results: the fact that transcriptions of imitations can be matched back to other category members (Category match questions) suggests that transcriptions still do carry some category information, so this is not a complete explanation of our results. Another possibility is that by selecting only the most frequent transcriptions, we unintentionally excluded less frequent transcriptions that were more diagnostic of category information.

Experiments 1 and 2 document a process of gradual change from an imitation of an environmental sound to a more word-like form. But do these emergent words function like other words in a language? In Experiment 3, we test the suitability of imitations taken from the beginning and end of transmission chains in serving as category labels in a category learning task.

## Experiment 3: Suitability of created words as category labels

If, as we claim, repeated imitation leads to more word-like forms, they should make for better category labels. For example, an imitation from a later generation may be easier to learn as a label for the category of sounds that motivated it than an earlier imitation, which is more closely yoked to a particular environmental sound. To the extent that repeating imitations abstract away the idiosyncrasies of a particular category member [38,39], it may also be easier to generalize later imitations to new category members. We tested these predictions using a category learning task in which participants learned novel labels for the categories of environmental sounds. The novel labels were transcriptions of either first or last generation imitations gathered in Experiment 1.

### Methods

#### Selecting words to learn as category labels

Of the 1814 unique words created through the transmission chain and transcription procedures, we sampled 56 words transcribed from first and last generation imitations that were equated in terms of length and match accuracy to the original sounds (see Supplementary Materials for additional details).

#### Procedure

Participants (*N*=67) were University of Wisconsin undergraduates. Participants were tasked with learning to associate novel labels (transcriptions of seed sounds) with the original seed sounds. Full instructions are provided in the Supplementary Materials. Participants were assigned between-subject to learn labels of either first or last generation imitations. On each trial, participants heard one of the 16 seed sounds. After a 1s delay, participants saw a label (one of the transcribed imitations) and responded *yes* or *no* using a gamepad controller depending on whether the sound and the word went together. Participants received accuracy feedback (a bell sound and a green checkmark if correct; a buzzing sound and a red “X” if incorrect). Four outlier participants were excluded due to high error rates and slow RTs.

Participants categorized all 16 seed sounds over the course of the experiment, but they learned them in blocks of 4 sounds at a time. Within each block of 24 trials, participants heard the same four sounds and the same four words multiple times, with a 50% probability of the sound matching the word on any given trial. At the start of a new block of trials, participants heard four new sounds they had not heard before, and had to learn to associate these new sounds with the words they had learned in the previous blocks.

### Results

Participants began by learning through trial-and-error to associate four written labels with four categories of environmental sounds. The small number of categories made this an easy task (mean accuracy after the first block of 24 trials was 81%; Fig. S5). Participants learning transcriptions of first or last generation imitations did not differ in overall accuracy, *p* = 0.887, or reaction time, *p* = 0.616.

After this initial learning phase (i.e. after the first block of trials), accuracy performance quickly reached ceiling and did not differ between groups *p* = 0.775. However, the response times of participants learning last generation transcriptions declined more rapidly with practice than participants learning first generation transcriptions, *b* = −114.13 (SE = 52.06), *t*(39.9) = −2.19, *p* = 0.034 (Fig. 4A). These faster responses suggest that, in addition to becoming more stable both in terms of acoustic and orthographic properties, repeated imitations become easier to process as category labels. We predict that given a harder task (i.e., more than four categories and 16 exemplars) would yield differences in initial learning rates as well.

Next, we examined specifically whether transcriptions from last generation imitations were easier to generalize to novel category exemplars by comparing RTs on trials immediately prior to the introduction of novel sounds (new category members) and the first trials after the block transition (±6 trials). The results revealed a reliable interaction between the generation of the transcribed imitation and the block transition, *b* = −110.77 (SE = 52.84), *t*(39.7) = −2.10, *p* = 0.042 (Fig. 4B). This result suggests that transcriptions from later generation imitations were easier to generalize to new category members.

**Figure 4.**
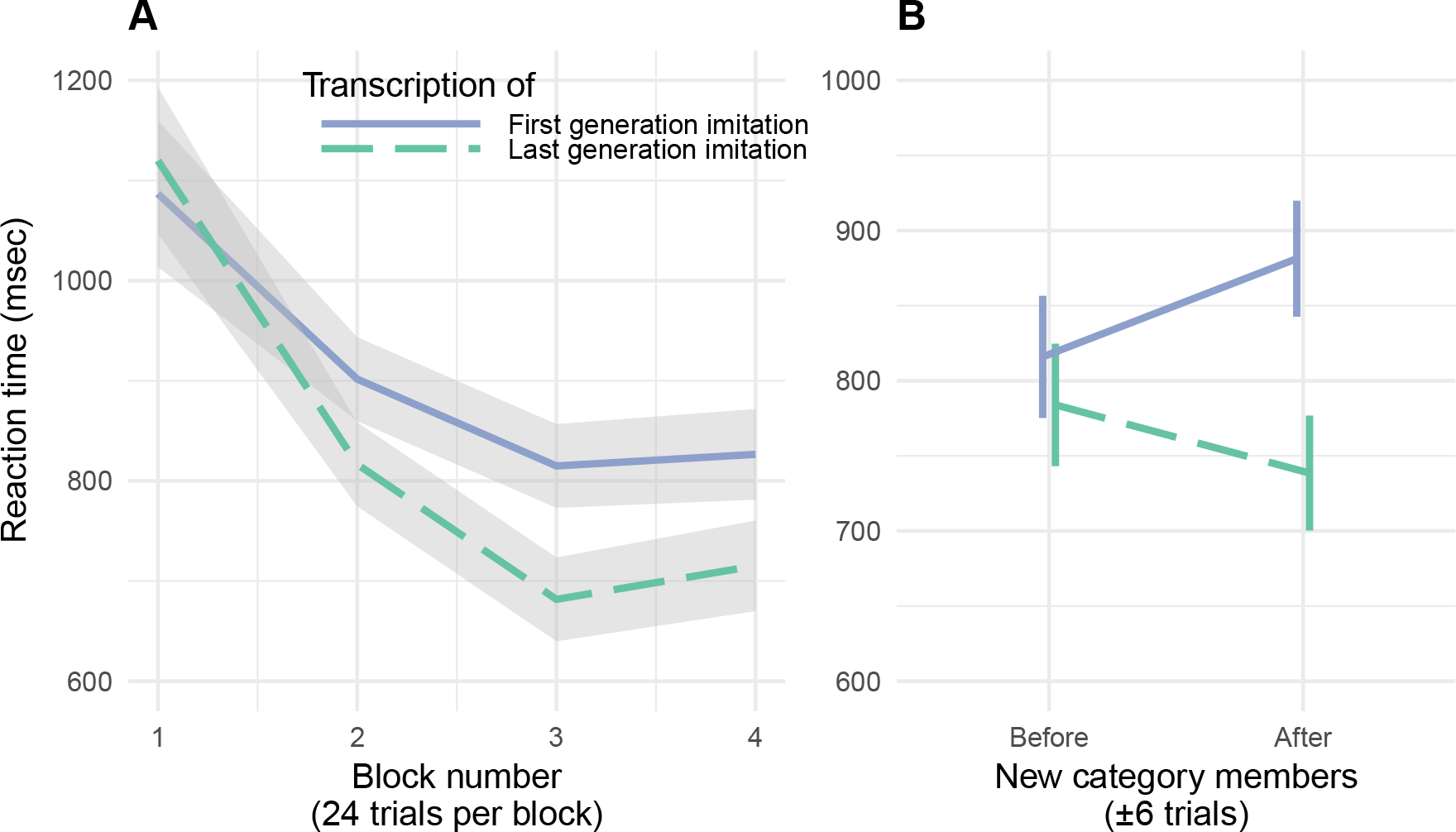
Repeated imitations made for better category labels. A. Mean RTs for correct responses in the category learning experiment with ±1 SE. B. Cost of generalizing to new category members with ±1 SE.

### Discussion

Transcriptions of vocal imitations that have undergone greater repetition were processed more quickly, and generalized to new category members more easily to new category members. These results point to a way that repeated imitation may lead to forms that become more word-like besides increase in stability.

## General Discussion

Accumulating evidence shows that iconic words are prevalent across the spoken languages of the world [23,24,30]. Counter to past assumptions about the limitations of human vocal imitation, people are surprisingly effective at using vocal imitation to represent and communicate about the sounds in their environment [33] and more abstract meanings [31]. These findings raise the possibility that early spoken words originated from vocal imitations, perhaps comparable to the way that many of the signs of signed languages appear to be formed originally from pantomimes [31,40]. Here, we examined whether simply repeating an imitation of an environmental sound — with no intention to create a new word or even to communicate — produces more word-like forms.

Our results show that through unguided repetition, imitative vocalizations became more word-like both in form and function. In form, the vocalizations gradually stabilized over generations, becoming more similar from imitation to imitation. The standardization was also found when the vocalizations were transcribed into English letters. Even as the vocalizations became more word-like, they maintained a resemblance to the original environmental sounds that motivated them. Notably, this resemblance appeared more resilient with respect to the category of sound (e.g., water-splashing sounds), rather than to the specific exemplar (a particular water-splashing sound). After eight generations the vocalizations could no longer be matched to the specific sound from which they originated any more accurately than they could be matched to the general category of environmental sound. Thus, information that distinguished an imitation from other sound categories was more resistant to transmission decay than exemplar information within a category. The resemblance to the original sounds was maintained even when the vocalizations were transcribed into a written form: participants were able to match the transcribed vocalizations to the original sound category at levels above chance.

We further tested the hypothesis that repeated imitation led to vocalizations becoming more word-like by testing the ease with which people learned the (transcribed) vocalizations as category labels (e.g., “pshfft” from generation 1 vs. “shewp” from generation 8 as labels for tearing sounds) (Exp. 3). Labels from the last generation were responded to more quickly than labels from the first generation. More importantly the labels from the last generation generalized better to novel category members. This fits with previous research showing that the relatively arbitrary forms that are typical of words (e.g. “dog”) makes them better suited to function as category labels compared to direct auditory cues (e.g., the sound of a dog bark) [38,39,41].

Compared to the large number of iconic signs in signed languages [8], the number of iconic words in spoken languages may appear to be very small [42,43]. However, increasing evidence from disparate language suggests that vocal imitation is, in fact, a widespread source of vocabulary. Cross-linguistic surveys indicate that onomatopoeia—iconic words used to represent sounds—are a universal lexical category found across the world’s languages [44]. Even English, a language that has been characterized as relatively limited in iconic vocabulary [45], is documented as having hundreds of onomatopoeic words not only for animal and human vocalizations (“meow”, “tweet”, “slurp”, “babble”, murmur”), but also for a variety of environmental sounds (e.g., “ping”, “click”, “plop”) [34,46]. Besides words that directly resemble sounds — the focus of the present study — many languages contain semantically broader inventories of ideophones. These words comprise a grammatically and phonologically distinct class of words that are used to express various sensory-rich meanings, such as qualities related to manner of motion, visual properties, textures and touch, inner feelings and cognitive states [44,47,48]. As with onomatopoeia, ideophones are often recognized by naïve listeners as bearing a degree of resemblance to their meaning [49].

Our study focused on imitations of environmental sounds as a source domain of meaning. Additional work is required to determine the extent to which vocal imitation can ground *de novo* vocabulary in other semantic domains [31,50]. Our hypothesis that vocal imitation may have played a role in the origin of some of the first spoken words does not preclude that gesture played an equal or more important role in establishing the first linguistic conventions [8,9,51]. In addition, the present studies—like nearly all experimental investigations of the evolution of language—are limited in their inferential power by the use of participants who already speak at least one language. It may turn out that the ability to repeat vocal imitations and converge on more word-like forms only arises in people who already know and use a full linguistic system, which would limit the relevance of our findings for the origins of spoken words.

Although our results show that repeated imitations lead to increases in stability of spoken (as well as transcribed) forms, we recognize that there are additional requirements for the vocalizations to be incorporated into a linguistic system. One of these may be familiarity with the referents that are being imitated. The extent to which our results depend on prior familiarity with the referents can be measured by extending our procedure to less familiar referential domains. Another design limitation is the use of auditory referents that can be imitated (environmental sounds). But although vocal imitation may seem to be restricted to auditory referents, prior results indicate that people show considerable agreement on how to vocally “imitate” non-auditory and even somewhat abstract meanings [31,50].

Among the qualities that distinguish natural language from other communication systems is the extreme diversity of signals (e.g. words) that individuals learn and use, and the speed with which these signals change over generations of speakers. As a consequence, the origins of most spoken words are opaque, making it difficult to investigate the process by which they were formed. Our experimental results show that the transition from vocal imitation to more word-like signals can, in some cases, be a rapid and simple process. The mere act of repeated imitation can drive vocalizations to become more word-like in both form and function with the vocalizations nevertheless still retaining some resemblance to their real-world referents. These findings suggest that repeated vocal imitation may constitute a significant mechanism for the origin of new words. It remains for future work to determine the extent to which the functioning of this process depends on the linguistic competencies of modern humans.

## Ethics

This was approved by the University of Wisconsin-Madison’s Educational and Social/Behavioral Sciences IRB and conducted in accordance with the principles expressed in the Declaration of Helsinki. Informed consent was obtained for all participants.

## Data, code, and materials

Our data, methods, materials, and analysis scripts, are available at osf.io/3navm.

## Authors’ contributions

P.E., M.P., and G.L. designed the research. P.E. conducted the research and analyzed the data. P.E., M.P., and G.L. wrote the manuscript.

## References

1. Seyfarth RM, Cheney DL. 1986 Vocal development in vervet monkeys. Animal Behaviour 34, 1640–1658.

2. Brysbaert M, Stevens M, Mandera P, Keuleers E. 2016 How Many Words Do We Know? Practical Estimates of Vocabulary Size Dependent on Word Definition, the Degree of Language Input and the Participant’s Age. Frontiers in Psychology 7, 55–11.

3. Wierzbicka A. 1996 Semantics: Primes and universals: Primes and universals. Oxford University Press, UK.

4. Evans N, Levinson SC. 2009 The myth of language universals: Language diversity and its importance for cognitive science. Brain and Behavioral Sciences 32, 429–492.

5. Pagel M, Atkinson QD, Meade A. 2007 Frequency of word-use predicts rates of lexical evolution throughout Indo-European history. Nature 449, 717–720.

6. Sapir E. 1921 Language: An introduction to the study of speech. New York: Harcourt, Brace; Company.

7. Labov W. 1972 Sociolinguistic patterns. University of Pennsylvania Press.

8. Goldin-Meadow S. 2016 What the hands can tell us about language emergence. Psychonomic Bulletin & Review 24, 1–6.

9. Kendon A. 2014 Semiotic diversity in utterance production and the concept of ‘language’. Philosophical Transactions of the Royal Society B: Biological Sciences 369, 20130293–20130293.

10. Klima ES, Bellugi U. 1980 The signs of language. Harvard University Press.

11. Frishberg N. 1975 Arbitrariness and Iconicity: Historical Change in American Sign Language. Language 51, 696–719.

12. Stokoe W. 1965 Dictionary of the American Sign Language based on scientific principles. Gallaudet College Press, Washington.

13. Wescott RW. 1971 Linguistic iconism. Linguistic Society of America 47, 416–428.

14. Goldin-Meadow S, Feldman H. 1977 The development of language-like communication without a language model. Science 197, 401–403.

15. Fay N, Lister CJ, Ellison TM, Goldin-Meadow S. 2014 Creating a communication system from scratch: Gesture beats vocalization hands down. Frontiers in Psychology 5, 663.

16. Arbib MA. 2012 How the brain got language: The mirror system hypothesis. Oxford University Press.

17. Armstrong DF, Wilcox S. 2007 The gestural origin of language. Oxford University Press.

18. Corballis MC. 2003 From hand to mouth: The origins of language. Princeton University Press.

19. Hewes GW. 1973 Primate Communication and the Gestural Origin of Language. Current Anthropology 14, 5–24.

20. Hockett CF. 1978 In search of Jove’s brow. American speech 53, 243–313.

21. Tomasello M. 2010 Origins of human communication. MIT press.

22. Pinker S, Jackendoff R. 2005 The faculty of language: what’s special about it? Cognition 95, 201–236.

23. Dingemanse M, Blasi DE, Lupyan G, Christiansen MH, Monaghan P. 2015 Arbitrariness, Iconicity, and Systematicity in Language. Trends in Cognitive Sciences 19, 603–615.

24. Perniss P, Thompson RL, Vigliocco G. 2010 Iconicity as a General Property of Language: Evidence from Spoken and Signed Languages. Frontiers in Psychology 1.

25. Clark HH, Gerrig RJ. 1990 Quotations as demonstrations. Language 66, 764–805.

26. Lewis J. 2009 As well as words: Congo Pygmy hunting, mimicry, and play. In The cradle of language, The cradle of language.

27. Brown RW, Black AH, Horowitz AE. 1955 Phonetic symbolism in natural languages. Journal of abnormal psychology 50, 388–393.

28. Dingemanse M. 2014 Making new ideophones in Siwu: Creative depiction in conversation. Pragmatics and Society

29. Donald M. 2016 Key cognitive preconditions for the evolution of language. Psychonomic Bulletin & Review, 1–5.

30. Imai M, Kita S. 2014 The sound symbolism bootstrapping hypothesis for language acquisition and language evolution. Philosophical Transactions of the Royal Society B: Biological Sciences 369.

31. Perlman M, Dale R, Lupyan G. 2015 Iconicity can ground the creation of vocal symbols. Royal Society Open Science 2, 150152–16.

32. Lemaitre G, Rocchesso D. 2014 On the effectiveness of vocal imitations and verbal descriptions of sounds. The Journal of the Acoustical Society of America 135, 862–873.

33. Lemaitre G, Houix O, Voisin F, Misdariis N, Susini P. 2016 Vocal Imitations of Non-Vocal Sounds. PloS one 11, e0168167–28.

34. Rhodes R. 1994 Aural images. Sound symbolism, 276–292.

35. Tamariz M. 2017 Experimental Studies on the Cultural Evolution of Language. Annual Review of Linguistics 3, 389–407.

36. Kirby S, Cornish H, Smith K. 2008 Cumulative cultural evolution in the laboratory: an experimental approach to the origins of structure in human language. Proceedings of the National Academy of Sciences 105, 10681–10686.

37. Hall KC, Allen B, Fry M, Mackie S, McAuliffe M. 2016 Phonological CorpusTools. 14th Conference for Laboratory Phonology

38. Edmiston P, Lupyan G. 2015 What makes words special? Words as unmotivated cues. Cognition 143, 93–100.

39. Lupyan G, Thompson-Schill SL. 2012 The evocative power of words: Activation of concepts by verbal and nonverbal means. Journal of Experimental Psychology: General 141, 170–186.

40. Fay N, Ellison TM, Garrod S. 2014 Iconicity: From sign to system in human communication and language. Pragmatics and Cognition 22, 244–263.

41. Boutonnet B, Lupyan G. 2015 Words Jump-Start Vision: A Label Advantage in Object Recognition. Journal of Neuroscience 35, 9329–9335.

42. Crystal D. 1987 The Cambridge Encyclopedia of Language. Cambridge Univ Press.

43. Newmeyer FJ. 1992 Iconicity and generative grammar. Language

44. Dingemanse M. 2012 Advances in the Cross-Linguistic Study of Ideophones. Language and Linguistics Compass 6, 654–672.

45. Vigliocco G, Perniss P, Vinson D. 2014 Language as a multimodal phenomenon: implications for language learning, processing and evolution. Philosophical Transactions of the Royal Society B: Biological Sciences 369, 20130292–20130292.

46. Sobkowiak W. 1990 On the phonostatistics of English onomatopoeia. Studia Anglica Posnaniensia 23, 15–30.

47. Nuckolls JB. 1999 The case for sound symbolism. Annual Review of Anthropology 28, 225–252.

48. Voeltz FE, Kilian-Hatz C. 2001 Ideophones. John Benjamins Publishing.

49. Dingemanse M, Schuerman W, Reinisch E. 2016 What sound symbolism can and cannot do: Testing the iconicity of ideophones from five languages. Language 92.

50. Perlman M, Lupyan G. 2018 People Can Create Iconic Vocalizations to Communicate Various Meanings to Naïve Listeners. Scientific Reports

51. Fay N, Arbib MA, Garrod S. 2013 How to Bootstrap a Human Communication System. Cognitive Science 37, 1356–1367.

